# Comprehensive analysis of central carbon metabolism reveals multiple connections between nutrients, biosynthetic capacity, and cell morphology in *Escherichia coli*

**DOI:** 10.1101/191585

**Authors:** Corey S. Westfall, Petra Anne Levin

## Abstract

Bacterial morphology is a complex trait that is highly sensitive to changes in the environment. For heterotrophic organisms, such as *Escherichia coli*, increases in nutrient levels are frequently accompanied by several-fold increases in both size and growth rate. Despite the dramatic nature of these changes, how alterations in nutrient availability translate into changes in growth and morphology remains a largely open question. To understand the signaling networks coupling nutrient availability with size and shape, we examined the impact of deletions in the entirety of non-essential central carbon metabolic genes on *E. coli* growth rate and cell size. Our data reveal the presence of multiple metabolic nodes that play important yet distinctive roles in shaping the cell. Consistent with recent work from our lab and others, although both are sensitive to nutrient availability, size and growth rate vary independently. Cell width and length also appear to be independent phenomena, influenced by different aspects of central carbon metabolism. These findings highlight the diversity of factors that can impact cell morphology and provide a foundation for further studies.

**Author summary:** Often taken for granted, the shape of bacterial cells is a complex trait that is highly sensitive to environmental perturbations. Nutrients in particular, strongly impact bacterial morphology together with growth rate. The ubiquitous, rod-shaped bacteria *Escherichia coli* increases both length and width several fold upon a shift from nutrient poor to nutrient rich medium, a change accompanied by an equally dramatic increase in growth rate. Central carbon metabolism is an obvious site for the integration of nutrient dependent signals that dictate cell size and shape. To develop a clearer picture of the molecular mechanisms coupling nutrient assimilation with cell growth and morphology, we screened the entirety of nonessential carbon metabolic genes for their contribution to growth rate and cell shape. Our data reveal the presence of multiple regulatory circuits coordinating different metabolic pathways with specific aspects of cell growth and morphology. Together, these data firmly establish a role for central carbon metabolism as an environmentally sensitive sculptor of bacterial cells.

## Introduction

The behavior and physiology of single celled organisms is at the mercy of their environment. Nutrients, in particular, dramatically impact the growth rate, cell cycle, and morphology of bacteria. *Salmonella typhimurium, Escherichia coli,* and *Bacillus subtilis* cells cultured in nutrient rich conditions exhibit mass doubling times up to 6-fold faster than their counterparts cultured in nutrient poor medium [1–3]. These increases in growth rate are accompanied by similarly dramatic increases in cell size. *E. coli* increases length (2-fold) and width (1.5-fold) for a ∼3-fold increase in total square area between nutrient poor and nutrient rich conditions [3]. B. *subtilis* increases length ∼3-fold in nutrient rich conditions while width remains more or less constant [2,4].

The positive relationship between nutrient availability, growth rate and cell size is generally referred to as the “Growth Law”. Besides cell size, DNA, RNA, and protein content of the cell also show a positive relationship with growth rate, with faster growing cells containing more of these macromolecules [1]. Although there is more total DNA/RNA/protein in larger, faster growing cells, their concentration remains relatively constant due to the concomitant increase in cell size [1,5].

Despite its name, the relationship between growth rate and cell size is complicated and there are several, well known exceptions to the Growth Law. Reducing growth rate by lowering culture temperature does not impact cell length or width, suggesting that it is nutrients rather than growth rate that determine cell size [6]. Conversely, mutations that lead to modest delays in division frequently result in increases in cell length without detectably impacting mass doubling time [7–9].

### Central carbon metabolism: a hub connecting nutrient availability with growth rate and cell size

As the entry and processing point for the raw materials required for cell growth and proliferation, central carbon metabolism (CCM) plays a key role in coordinating nutrient availability with cell growth and size. Recent work indicates that CCM impacts bacterial morphology in several ways: by its impact on cell cycle progression [10–12]; by its impact on lipid synthesis [6]; and through its impact on cell wall biogenesis [13].

CCM-mediated changes in cell cycle progression primarily affect cell length, while changes in lipid synthesis and cell wall biogenesis impact both length and width. In an example of the former, UDP-glucose, produced in two steps from glucose-6-phosphate at the beginning of glycolysis, stimulates interactions between the moonlighting glucosyltransferase OpgH in *E. coli* and the unrelated enzyme UgtP in B. *subtilis* and the essential tubulin-like cell division protein FtsZ, delaying division and increasing cell length during growth in carbon rich medium [4,12]. In contrast, defects in fatty acid synthesis (FAS) lead to reductions in both length and width, effectively recapitulating the impact of nutrient limitation on cell morphology [6,14].

CCM can also effect morphology via its impact on the availability of cellular building blocks, particularly those required for synthesis of the cell wall. In a recent study, the Young laboratory determined that defects in *wecE*, a gene encoding an aminotransferase important for making the *E. coli* outer membrane molecule enterobacterial common antigen (ECA), led to the formation of swollen, filamentous cells. ECA is transported to the cell wall via undecaprenyl-phosphate, the lipid carrier responsible for shuttling precursors for peptidoglycan and other components of the bacterial cell envelope across the plasma membrane. Careful genetics revealed the morphological defects associated with defects in *wecE* were due to sequestration of undecaprenyl-phosphate conjugated to an ECA intermediate, interfering with peptidoglycan synthesis [15].

More generically, nutrient starvation leads to an accumulation of the small molecule guanosine tetraphosphate (ppGpp) that interacts directly with RNA polymerase, to inhibit transcription of a wide array of genes, effectively curtailing flux through multiple biosynthetic pathways [16]. ppGpp concentration is inversely proportional to both size and growth rate: ectopic expression of the ppGpp synthetase RelA during growth in nutrient rich medium reduces both growth rate and cell size [6,17]. The reduction in size is indirect and dependent on ppGpp-mediated inhibition of fatty acid synthesis, while the reduction in growth rate is almost certainly a consequence of ppGpp-dependent inhibition of a host of essential processes [6].

While these examples highlight several ways in which CCM might impact the positive relationship between cell growth and cell morphology, they represent only a small fraction of the pathways that constitute CCM. To identify additional connections between nutrient levels, metabolism, and cell size in *E. coli*, we screened non-essential (CCM) gene deletions for defects in cell size and growth rate. Our data highlight three CCM nodes—the first branch-point of glycolysis, the pentose-phosphate pathway, and acetyl-CoA metabolism—as critical links between nutrient availability, cell growth and cell size. While some aspects of CCM bottlenecks appear to directly impact the relationship between nutrient availability and biosynthetic flux, other aspects of CCM effect growth and morphology in more nuanced ways.

## Results

### Multiple connections exist between CCM, growth rate, and cell size

To identify key steps of CCM that impact the growth and morphology of *E. coli*, we phenotypically analyzed cells defective in 44 non-essential CCM genes (S1 Table) to. These experiments took advantage of the Keio Collection, an ordered insertion-deletion library of all non-essential genes in *E. coli* [18] and Coli-Inspector, an Image J plug in designed for high throughput analysis of bacterial morphology [19]. Prior to analysis, the Keio insertion-deletions were transduced into the laboratory wild type strain MG1655. Briefly, single colonies of each mutant were picked and cultured under nutrient rich conditions (Luria-Bertani (LB) broth supplemented with 0.2% glucose) to an OD_600_ of ∼0.2. Cultures were then back-diluted to OD_600_ of 0.01 and tracked for 4 generations until they reached a maximum OD_600_ of ∼0.2 (the low OD_600_ ensures cells are all actively growing and at approximately the same growth phase prior to analysis). Cells were sampled, fixed, and imaged by phase contrast microscopy. Cell size (length and width) was determined using Coli-Inspector in ImageJ.

Based on size and mass doubling time, we assigned the 44 mutants into six classes (Table 1). Represented by two mutations, Class I cells were >10% smaller than wild type cells with mild to undetectable changes in growth rate (within two SD of wild type mass doubling time). Eight mutations fell into Class II: small (more than 10% decrease in square area than wild type) with a significant decrease in doubling time (>20% wild type). Class III was heterogenous, consisting of a population dominated by smaller than normal cells and punctuated by the presence of a few (5-10%) very long cells. Three mutants fell into Class IV long (>10% longer than WT) and two into Class V, heterogenous with a wide distribution of cell lengths and variable cell widths (Fig 1A). Surprisingly, the majority of mutants (26) were essentially wild type with regard to growth rate and cell size (Class VI). This finding suggests the presence of alternative pathways that permit *E. coli* to bypass particular CCM enzymes and/or that nutrients that allow for bypass are present in LB. We elected not to investigate the impact of deletions in either ppsA, the first step of gluconeogenesis, or *fumC*, one of the three isoforms involved in the conversion of fumarate to malate in the TCA cycle, on cell size as we were unable to obtain stable MG1655 transductants of either mutant. Representative images of each mutant are shown (Fig 2).

**Table 1:**
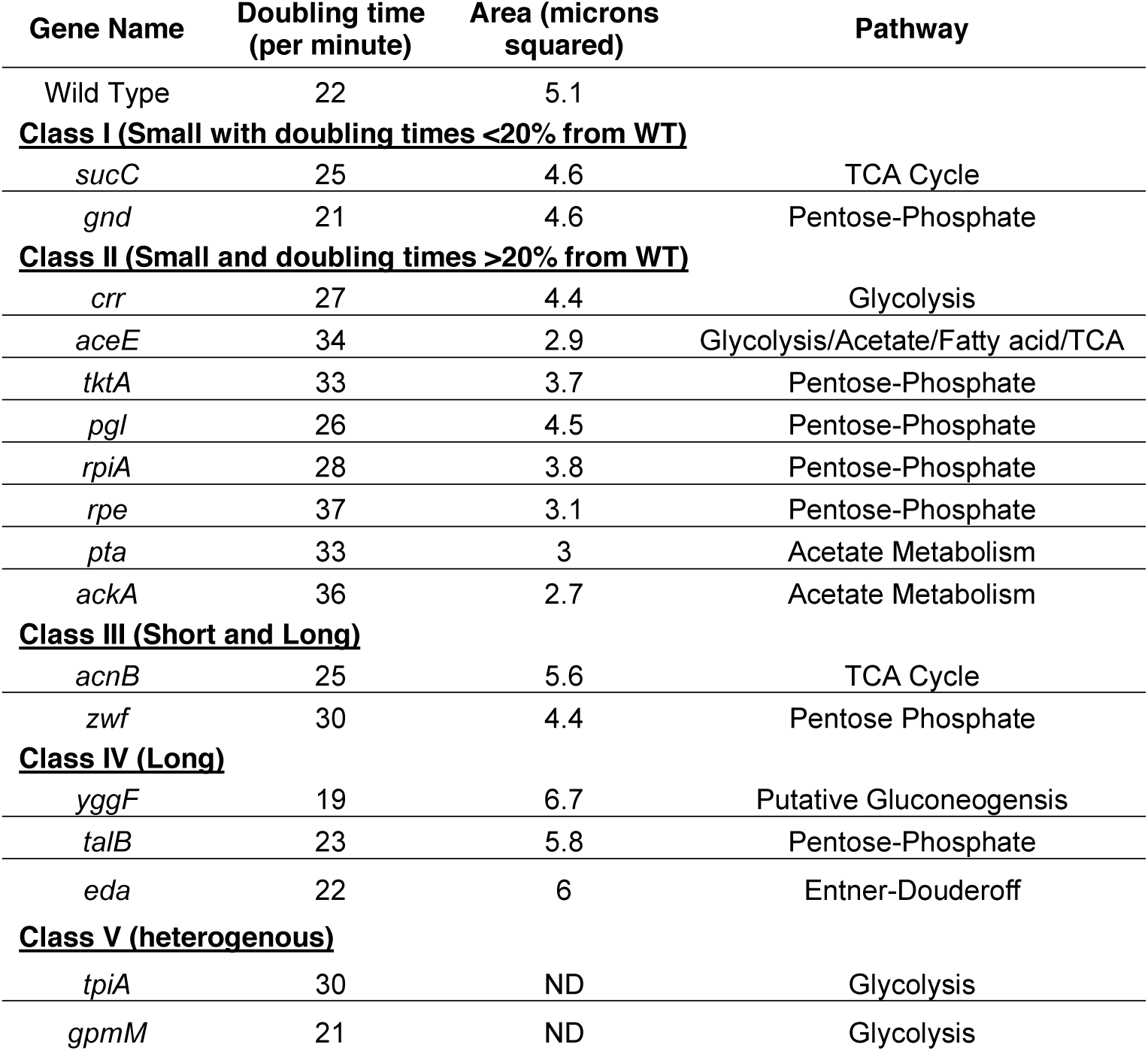
CCM deletion mutants exhibiting significant alterations in growth rate and/or morphology.

| Gene Name | Doubling time<br>(per minute) | Area (microns<br>squared) | Pathway |
| --- | --- | --- | --- |
| Wild Type | 22 | 5.1 |  |
| **<u>Class I (Small with doubling times <20% from WT)</u>** |  |  |  |
| *sucC* | 25 | 4.6 | TCA Cycle |
| *gnd* | 21 | 4.6 | Pentose-Phosphate |
| **<u>Class II (Small and doubling times >20% from WT)</u>** |  |  |  |
| *crr* | 27 | 4.4 | Glycolysis |
| *aceE* | 34 | 2.9 | Glycolysis/Acetate/Fatty acid/TCA |
| *tktA* | 33 | 3.7 | Pentose-Phosphate |
| *pgl* | 26 | 4.5 | Pentose-Phosphate |
| *rpiA* | 28 | 3.8 | Pentose-Phosphate |
| *rpe* | 37 | 3.1 | Pentose-Phosphate |
| *pta* | 33 | 3 | Acetate Metabolism |
| *ackA* | 36 | 2.7 | Acetate Metabolism |
| **<u>Class III (Short and Long)</u>** |  |  |  |
| *acnB* | 25 | 5.6 | TCA Cycle |
| *zwf* | 30 | 4.4 | Pentose Phosphate |
| **<u>Class IV (Long)</u>** |  |  |  |
| *yggF* | 19 | 6.7 | Putative Gluconeogenesis |
| *talB* | 23 | 5.8 | Pentose-Phosphate |
| *eda* | 22 | 6 | Entner-Doudoroff |
| **<u>Class V (heterogenous)</u>** |  |  |  |
| *tpiA* | 30 | ND | Glycolysis |
| *gpmM* | 21 | ND | Glycolysis |

**Figure 1:**
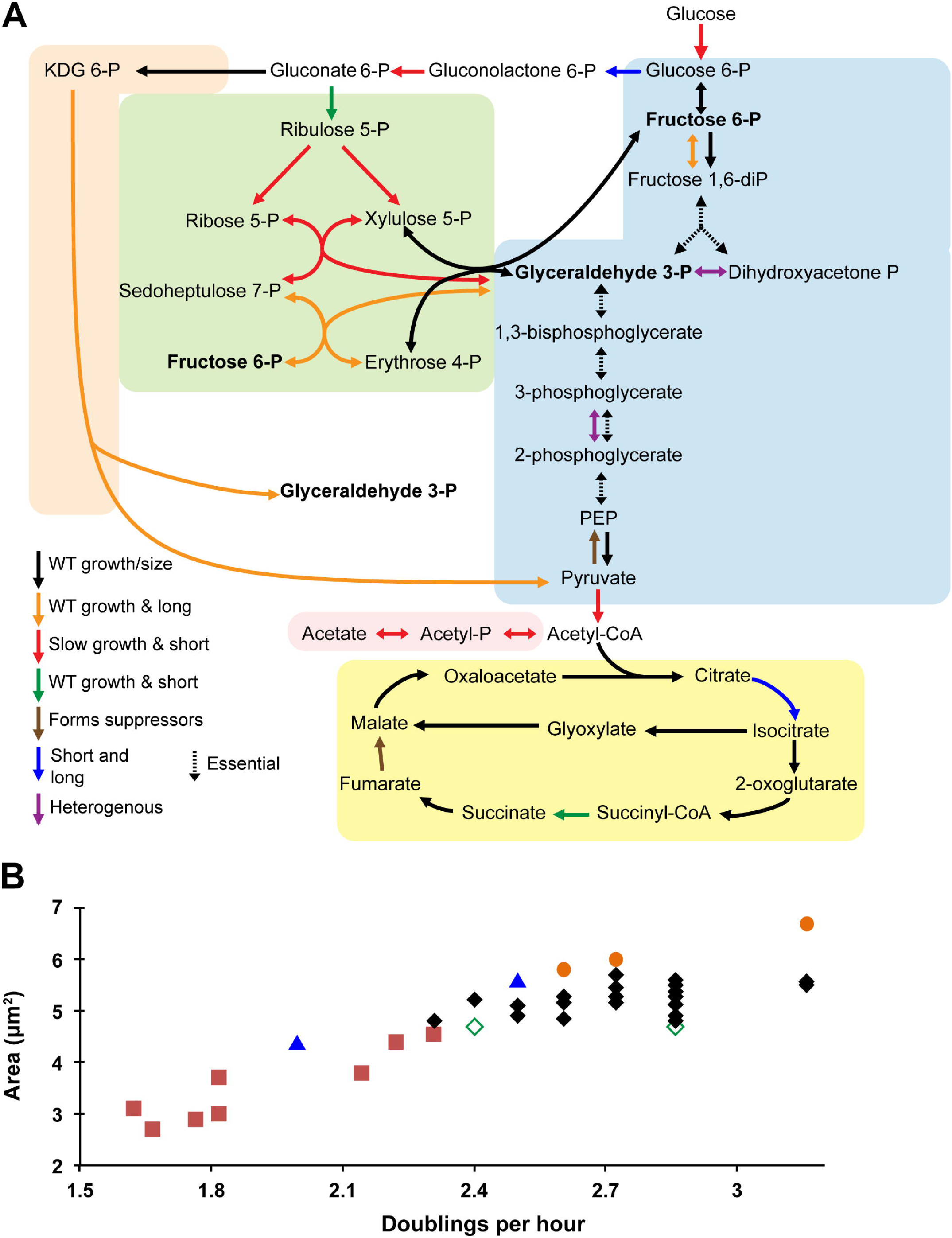
The effect of central carbon metabolic gene deletions on cell growth and size. (A) Mutant phenotypes mapped onto central carbon metabolic pathways. Arrows are colored based on the phenotype as follows: black= normal growth and size; green = normal growth and small size (<10% smaller than WT); red = slow growth (>20% increase in doubling time) and small size; orange = normal growth and long cells (>10% larger than WT); blue = short and long cells; purple = highly variable size; and dotted lines = essential genes. Brown arrows correspond to deletions that obtained suppressor mutations rapidly and were not analyzed. (B) Relationship between cell area and growth rate, as doublings per hour. Icons are colored as in panel (A).

**Figure 2:**
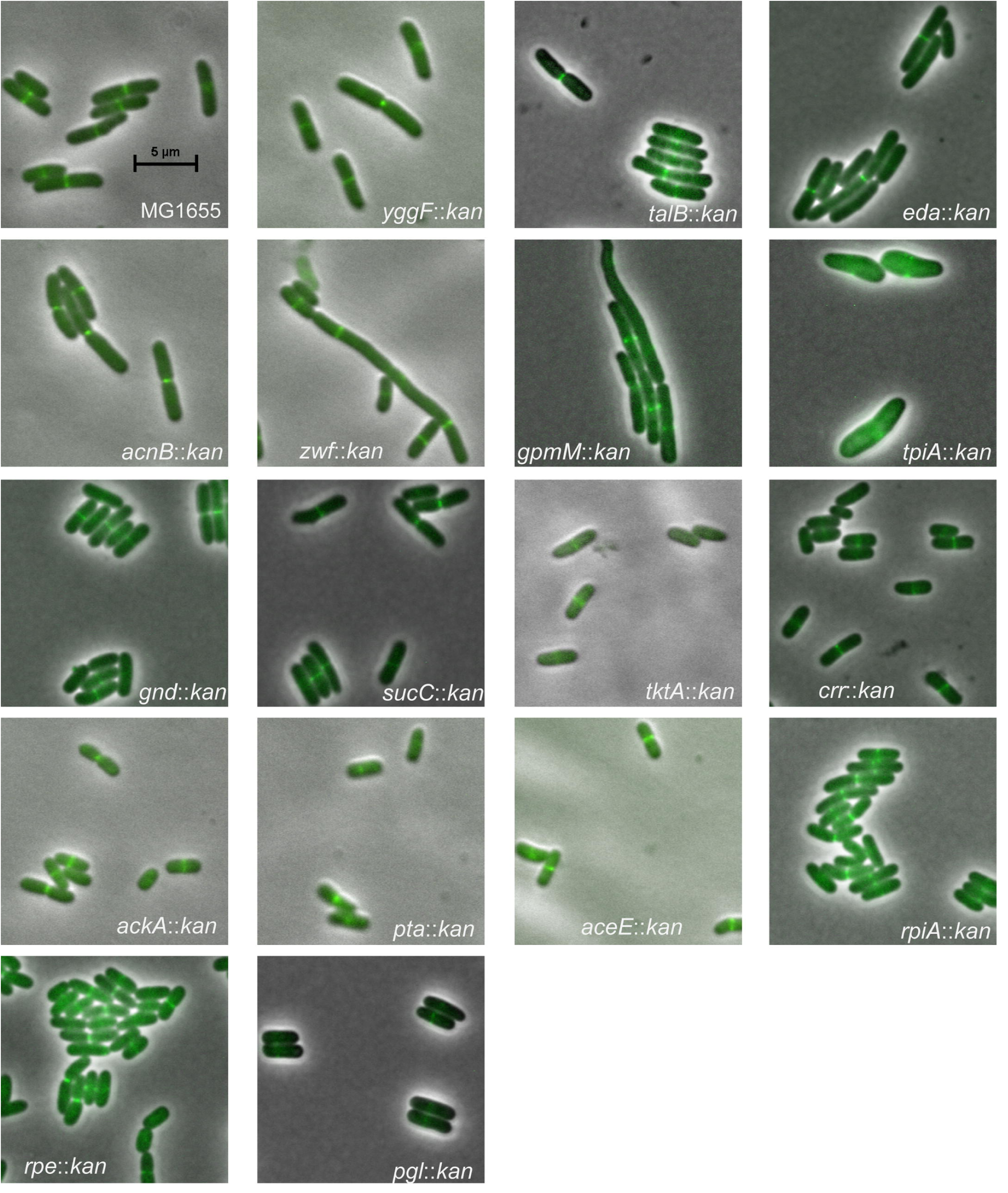
Representative images of all CCM mutants exhibiting morphology distinct from the wild type parental strain. Merged images of morphologically distinct CCM mutant strains expressing an *ftsZ-GFP* fusion to permit visualization of the cytokinetic ring. Scale bar = 5 microns and is the same for all images.

Of the different classes, only the eight Class II mutants “obeyed” the Growth Law, with reductions in growth rate accompanied by predictable reductions in cell size (Fig 1B). The close correlation between growth rate and size for Class II mutants suggests that the loss of individual genes and/or their product’s catalytic function, leads to a biosynthetic bottleneck that recapitulates the impact of nutrient starvation on cellular physiology. In contrast, despite their heterogeneity with regard to cell length, both Class I and Class IV mutants exhibited more or less normal growth rates, suggesting a potential defect in the signal transduction pathways coordinating CCM with cell growth and cell size. The heterogeneity of Class III and Class V indicate a loss of proper regulation of cell cycle progression and/or width control, leading to stochastic variations in cell morphology.

### Growth in carbon poor medium largely ameliorates CCM mutant morphological defects

To clarify the impact of nutrient flux on individual CCM mutant phenotypes, we assessed the growth rate and size of Class I-V mutants in three different media: LB + 0.2% glucose, the minimal media AB + 0.2% glucose, and AB + 0.2% succinate representing high nutrient conditions, low nutrient glycolytic conditions, and low nutrient gluconeogenic conditions respectively. Cells were measured using the Matlab software package SuperSegger, which facilitates the rapid analysis of large numbers of clumped cells [20]. In general, SuperSegger results corresponded almost exactly with the Coli-Inspector data reported above, although cell areas were smaller over all most likely due to a difference in how the two programs define cell boundaries and at what point of the cell cycle that they “divide” the parent cell into two daughter cells (see wild type area in Table 1 and S3 Table).

In light of data indicating that cells defective in *crr,* encoding the phosphotransferase system subunit EIIA^glc^, exhibited significant reductions in size (Table 1), we also examined the nutrient dependent phenotype of cells defective for adenylate cyclase or the cAMP receptor (Crp) production. Crr is part of a phosphodependent transferase system that is essential for the import of glucose and other sugars into *E. coli* [21]. Crr facilitates glucose import via transfer of a phosphate group from Hpr to EIIB^glc^ and finally to glucose, generating glucose-6-phosphate at the beginning of glycolysis (reviewed in [22]). During growth on a non-glycolytic substrate, phosphorylated EIIA^glc^ accumulates and activates the adenylate cyclase, CyaA, leading to synthesis of cyclic-AMP (cAMP) [23,24]. A coactivator, cAMP binds to Crp facilitating DNA binding and stimulating transcription of a broad swathe of genes including those involved in the transport and utilization of non-glycolytic carbon sources [25,26]. If *crr* impacts cell size via its impact on cAMP production and Crp activation, defects in both *cyaA* and *crp* should be phenotypically identical to *crr* mutants with regard to cell size and growth rate. We found that this is indeed true as deletions in *crr, cyaA*, or *crp* led to similar growth rates (29, 29, and 28 minutes) and cell size (2.1 square micron area for all three mutants). This implies that the growth rate and size phenotypes of *crr::kan* are caused not by a lack of glucose import but instead by blocking cAMP synthesis.

Generally speaking, most of the CCM mutants retained a positive relationship between nutrient availability and cell size with the exception of the three cAMP mutants, *crr::kan, cyaA::kan,* and *crp::kan* (Fig 3). During growth in carbon poor and/or gluconeogenic conditions, the lengths of Class I and Class II mutants approached that of wild type, particularly in AB-succinate medium, and the heterogeneous phenotype exhibited by mutants in Class III and Class IV was largely relieved. Width also decreased, although more modestly. The average width of the mutants ranged from 0.68-0.86 microns during growth in LB-glu and 0.48-0.60 microns during growth in AB-suc. Consistent with catabolite repression serving to inhibit division during growth on a gluconeogenic carbon source, the *crp*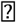*kan* mutants (dotted line in Fig 3B, 3C, and 3D) did not show any change in size when switched from LB-glu to AB-glu. Both *crr::kan* and *cyaA::kan* (magenta and gray in Fig 3) were similar to *crp::kan*, showing only minor, insignificant changes in length, with no change in width. Unfortunately, *crr, cyaA,* and *crp* are essential under gluconeogenic conditions, preventing analysis of these mutants in AB-succinate.

**Figure 3:**
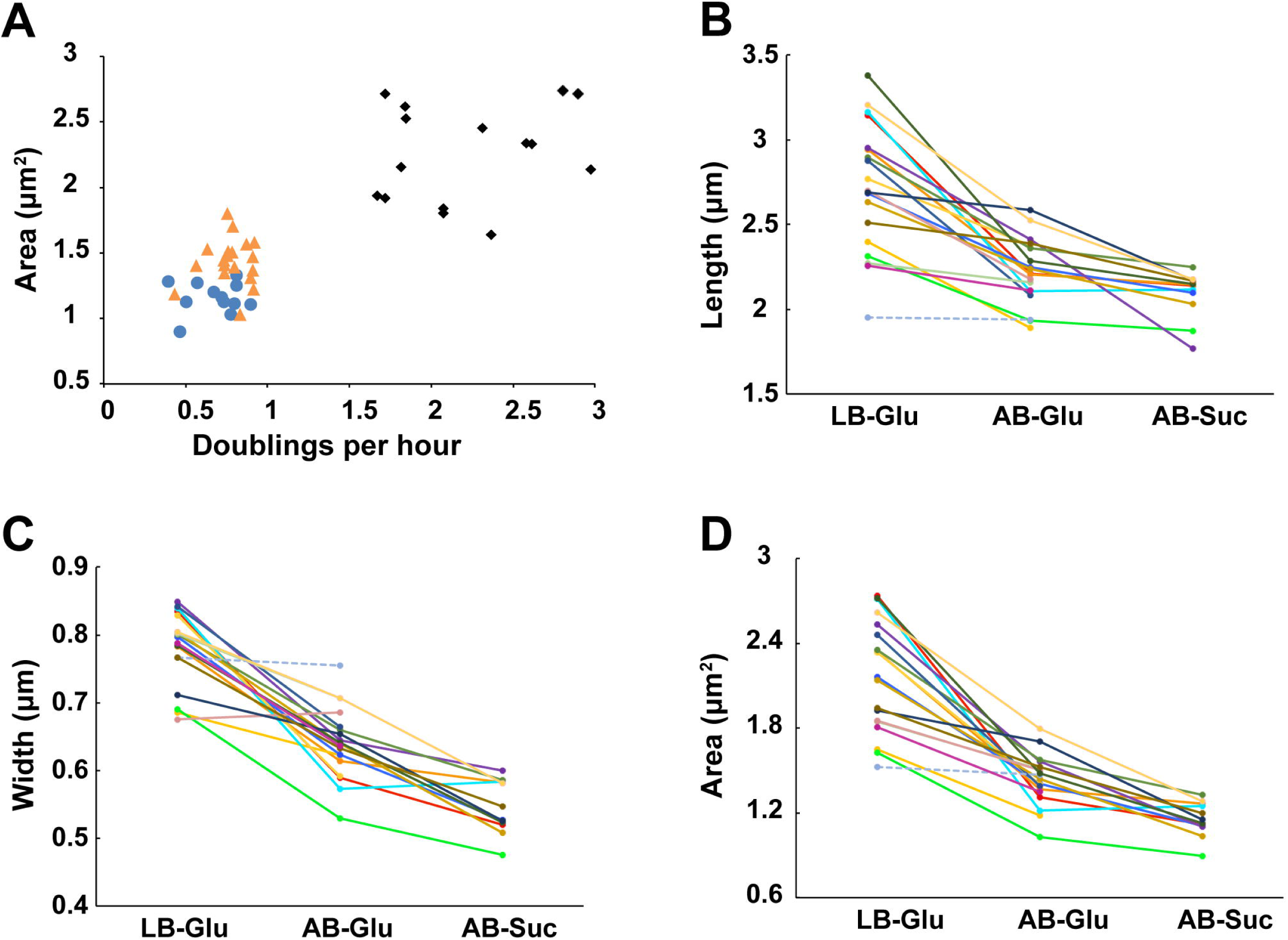
CCM deletions exhibit medium-dependent effects on growth and morphology. (A) Cell size versus growth rate in LB-glucose (black diamonds), AB-glucose (orange triangles), and AB-succinate (blue circles). Length (B), width (C), and area (D) are plotted for each mutant in the three different media. Dotted line = *crp::kan* in LB-glucose and AB-glucose.

Although we were able to obtain cell size data for *rpiA, crr, rpe, aceE*, and *sucA* in carbon rich (LB-glu) and carbon poor (AB-glu) conditions, all five mutants were unable to grow in the gluconeogenic condition, AB + 0.2% succinate. At first glance this finding appears to be in conflict with data indicating that *rpiA, rpe, aceE*, and *sucA* are capable of growth in another carbon poor gluconeogenic condition: M9 + 0.1% glycerol [27]. However, this discrepancy likely reflects the ability of glycerol to enter the CCM partway through glycolysis—as dihydroxyacetone phosphate—while succinate enters through the TCA cycle, leading to dependence on different pathways through CCM to synthesize all needed carbon metabolites.

### Defects in CCM differentially impact assembly of the cell division machinery

To gain insight into the relationship between growth rate, cell cycle progression and CCM during growth in LB-glucose, we assessed the ability of different mutants to suppress the heat sensitivity associated with the GTP binding defective FtsZ mutant, *ftsZ84* (G105S) [28]. While *ftsZ84* supports both FtsZ assembly and division under permissive conditions (LB at 30°C), under nonpermissive conditions (LB with no salt at 42°C), FtsZ assembly and division are blocked, leading to filamentation and cell death [7,29]. Based on extensive work from our group and others, conditions or mutations that enhance FtsZ assembly are expected to suppress the heat sensitivity of temperature sensitive *ftsZ* alleles [4,17,30,31]. Changes in the availability of CCM products or components that normally inhibit FtsZ assembly should restore growth and colony formation to *ftsZ84* mutants under non-permissive conditions while defects in CCM-related factors that normally promote FtsZ assembly will enhance *ftsZ84* heat sensitivity. Double mutants in CCM deletions and ftsZ84 were generated to examine the impact of these mutants on FtsZ assembly.

Consistent with a positive impact on FtsZ assembly, the majority of Class I-V mutants promoted growth of *ftsZ84* cells under mildly restrictive conditions (37°C, LB no salt) (Fig 4A). The strongest suppressors were loss-of-function mutations in *aceE* or *ackA* both of which increased *ftsZ84* cell viability 10,000-fold during growth at 37°C on salt-free LB medium. Mild suppression was observed in loss of function mutations in *crr, tktA*, and *pta* which increased *ftsZ84* cell viability 1,000-fold, 100-fold, and 100-fold respectively. The *zwf* deletion was alone in leading to enhancement of *ftsZ84*, i.e. lack of growth at 30° on LB-no salt (Fig 4B). Combined with the short and long phenotype of *zwf*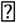*kan*, it appears that loss of *zwf* leads to impairment of FtsZ activity. Significantly, we did not observe a consistent correlation between reduced growth rate and suppression of *ftsZ84* heat-sensitivity among the various CCM mutants.

**Figure 4:**
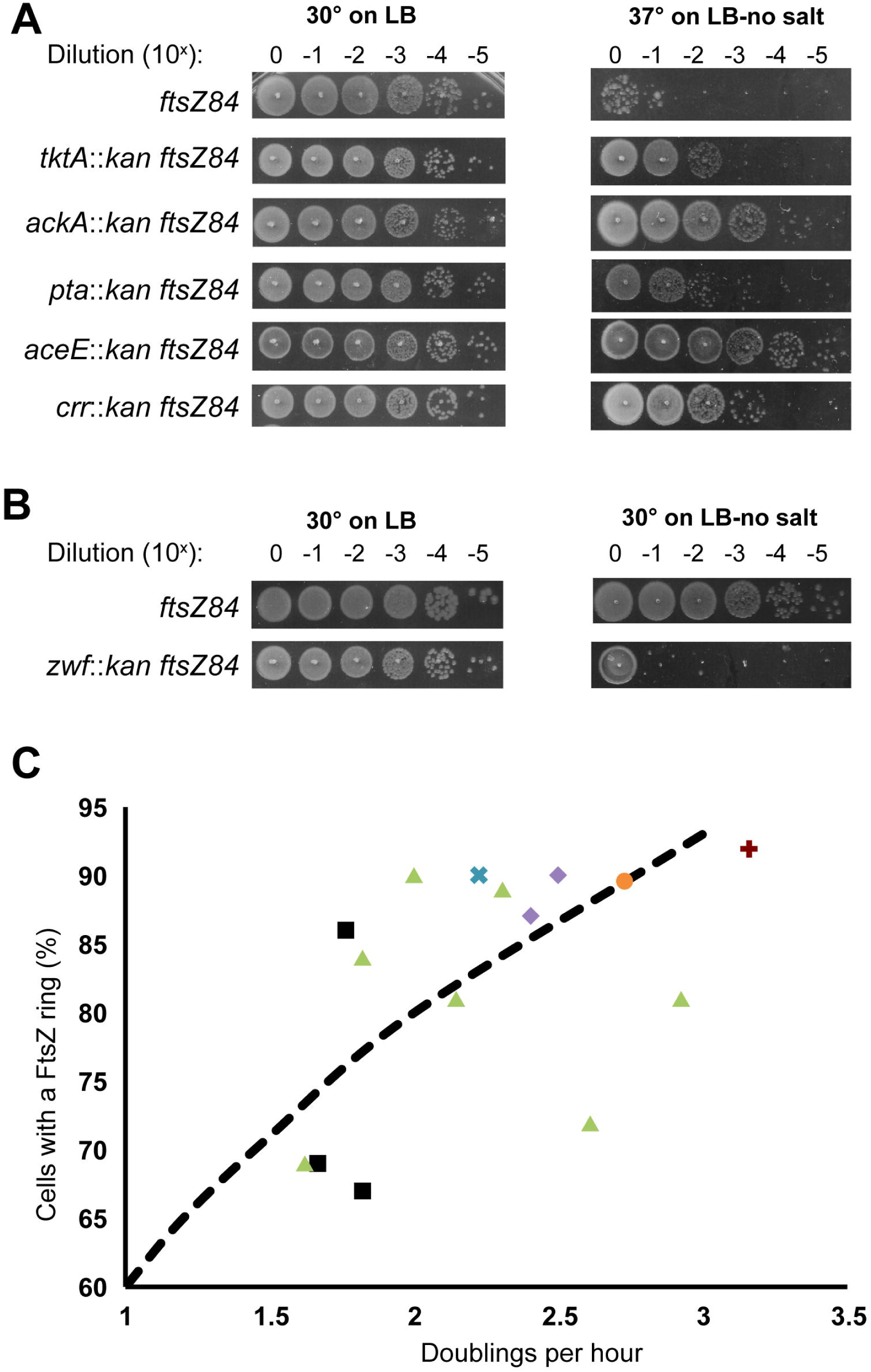
CCM mutations differentially impact FtsZ localization and activity. (A) Dot-plating efficiencies of the CCM deletion *ftsZ84* double mutants cultured under permissive (LB, 30°C) and non-permissive (LB-no salt, 37°C) conditions. (B) Dot-plating efficiency of *zwf::kan ftsZ84* double mutant. Note the reduction in *zwf::kan ftsZ84* viability under intermediate growth conditions (30°C on LB-no salt). (C) FtsZ ring frequency versus growth rate for the CCM mutants. For reference we have included previously reported data on the relationship between nutrient availability and FtsZ ring frequency (dotted black line) [12]. Mutants are grouped by metabolic pathway: black squares = acetyl-CoA production and acetate fermentation; green triangles = pentose-phosphate shunt; purple diamonds = TCA cycle; orange circle = Entner-Doudoroff pathway; blue cross = cAMP synthesis; and red plus = gluconeogenesis.

As part of these experiments we also revisited the impact of a loss of function mutation in *pgm*, encoding phosphoglucomutase—the enzyme that converts glucose-6-phosphate to glucose-1-phosphate. In previous work, we had reported that deletions in *pgm* suppress *ftsZ84* heat sensitivity [12]. However, we were unable to detect any suppression of *ftsZ84* by *pgm* in this instance. Resequencing BH173, the strain employed in the original study revealed acquisition of a reversion mutation that restored *ftsZ84* sequence to wild type.

To gain a better understanding of the mechanisms coordinating central carbon metabolism with cell morphology, we undertook an in depth analysis of two distinctive phenotypes: the reduction in length and width observed in cells defective in catabolite repression and the link between acetate metabolism, cell growth and cell cycle progression revealed by defects in *aceE, ptaA*, and *ackA* (Fig 7). The results of these efforts are described below.

### Catabolite-dependent regulation of FtsZ assembly and function

Reasoning that if the product of crr, EIIA^glc,^ was impacting division indirectly via its role in stimulation of adenylate cyclase and subsequent activation of the Crp transcription factor, we hypothesized that 1) defects in either the adenylate cyclase (encoded by *cya*) or *crp* should restore growth to *ftsZ84* cells under non-permissive conditions, and 2) the addition of extracellular cAMP (which is able to diffuse through the membrane [32]) to *crr::kan ftsZ84* cells should bypass *crr::kan* mediated suppression of *ftsZ84* heat sensitivity but only in the presence of Crp. Consistent with this model, deletion of either *cya* or *crp* phenocopied *crr*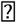*kan* mediated *ftsZ84* suppression at 37°C on salt free medium (Fig 5). More directly, 5 mM cAMP was sufficient to restore heat sensitivity to *crr::kan ftsZ84* cells as well as cells defective in adenylate cyclase (*cyaA*) but only in the presence of *crp* (Fig 5). Together, these data strongly argue for the presence of at least one inhibitor of FtsZ assembly within the extensive Crp regulon.

**Figure 5:**
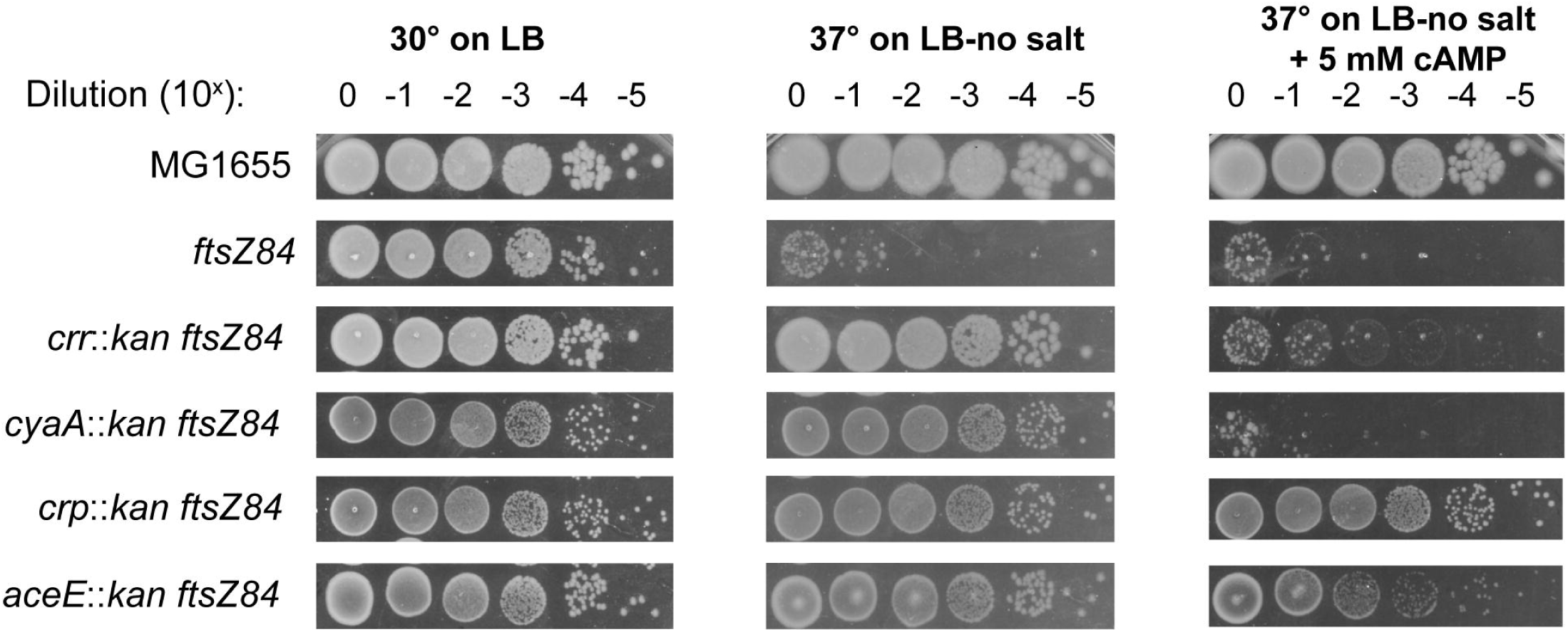
Loss of cAMP signaling suppresses ftsZ84 heat-sensitivity. Representative dot-plating efficiencies of *ftsZ84* alone or in combination with insertion deletions in cAMP signaling genes *crr, cyaA*, and *crp.* Cells were plated on and grown at permissive (LB, 30°C), non-permissive (LB-no salt, 37°C), and nonpermissive containing cAMP (LB-no salt + 5mM cAMP, 37°C) conditions.

### cAMP controls cell width through regulation of the morphogene *bolA*

In contrast to wild type cells, which decrease in both length and width upon a shift from glycolytic (low cAMP) to gluconeogenic (high cAMP) conditions, the *crp* deletion showed no significant changes in width when changing media from LB-glu to AB-glu, suggesting a potential role for cAMP in cell width regulation (Fig 3). To explore this idea, we examined the impact of extracellular cAMP on wild type (MG1655) cells cultured in both glycolytic and gluconeogenic conditions.

Consistent with a negative impact on cell width, 5 mM exogenous cAMP was sufficient to decrease wild type cell width in both LB and AB + 0.2% glucose (0.96 to 0.81 microns and 0.73 to 0.68 microns respectively) (Fig 6A). The addition of cAMP also significantly decreased cell length (3.8 to 3.2 microns for LB and 2.7 to 2.5 microns for AB-Glu) and growth rate (2.7 to 2.3 doubling per hour for LB and 0.98 to 0.88 doublings per hour for AB-Glu) (Fig 6B). Significantly, cAMP-dependent reductions in size are independent of growth rate. Reducing levels of exogenous cAMP from 5 mM to 1 mM decreased the growth rate of *cyaA::kan* (which is unable to synthesize any cAMP) in gluconeogenic conditions (AB + 0.2% succinate) from 0.87 to 0.79 doublings per hour, while cell width increased from 0.65 to 0.75 microns (Fig 6C). Although width was independent of growth rate, length was not as the *cyaA::kan* mutant length decreased with lower cAMP from 2.6 microns in 5 mM cAMP to 2.2 microns in 1 mM cAMP in a manner similar to the growth rate (Fig 6D).

**Figure 6:**
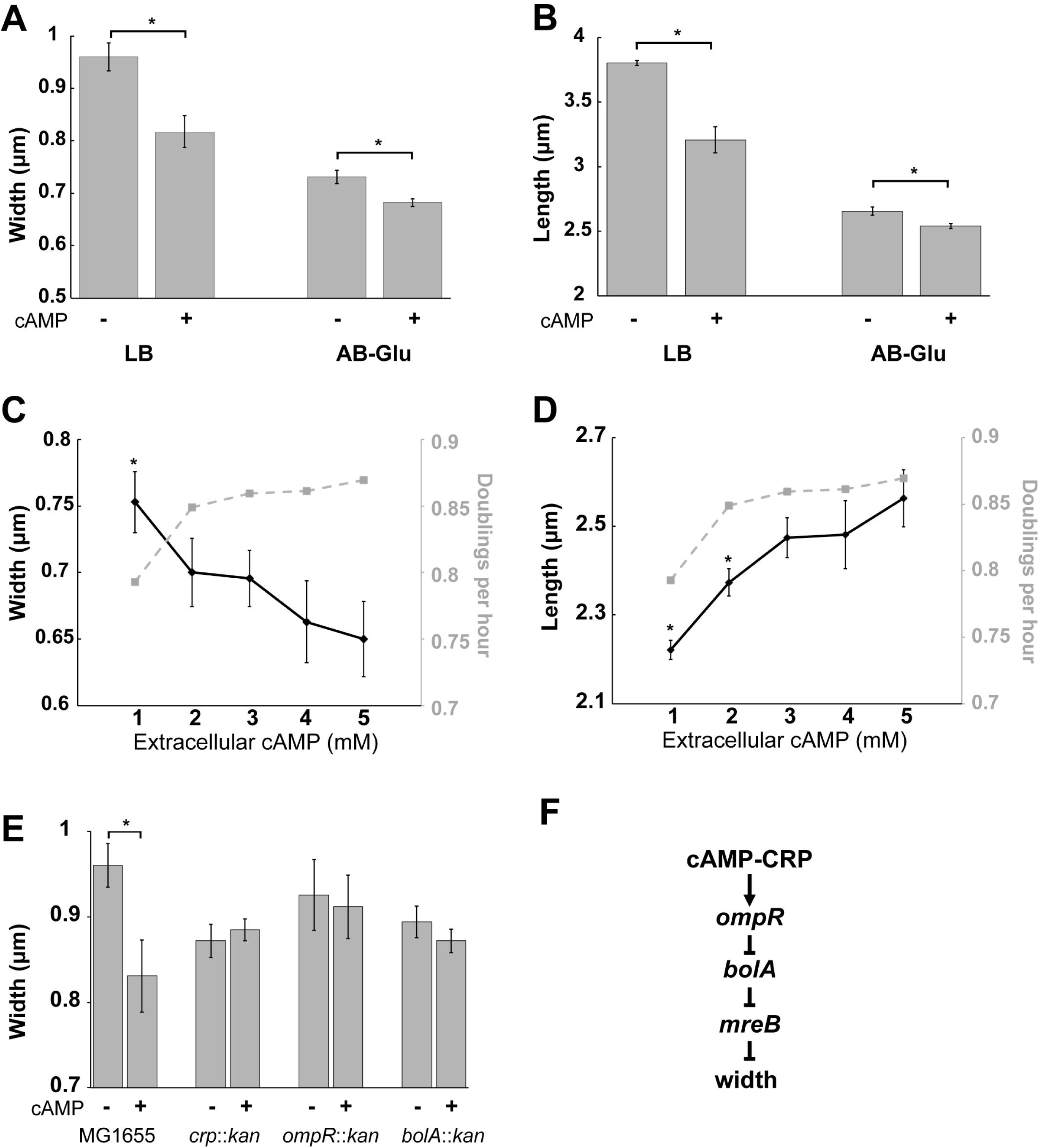
cAMP signaling reduces cell width via the morphogenic transcription factor, *bolA.* Wild-type MG1655 cell size was measured with and without exogenous 5 mM cAMP in either LB or AB-glucose. Width is shown in (A) and length in (B). Width (C) and length (D) of *cyaA::kan* grown in AB-succinate with different levels of exogenous cAMP shows that width is not correlated with growth rate but length is. (E) Deletions in *crp, ompR*, or *bolA* abolish the cAMP-dependent cell reductions in cell width. (F) Proposed genetic pathway linking cAMP to cell width.

cAMP has been implicated in the Crp dependent activation of the transcription factor OmpR, which inhibits expression of a host of genes, including several whose products have morphogenic activity [33]. Of particular interest is *bolA*, encoding a transcription factor that inhibits transcription of *mreB* [34], which encodes a key component of the elongation machinery [35]. Reductions in MreB activity lead to a block in synthesis of lateral cell wall material and the formation of wide and rounded cells [36,37]. *bolA* transcription has previously been shown to be inhibited by addition of extracellular cAMP [38].

Together, genetic interactions between *crp, ompR*, and *bolA*, support a model in which cAMP increases cell width indirectly, via down regulation of OmpR and the concomitant increase in BolA expression leading to repression of MreB mediated cell wall synthesis. Consistent with this model, cells defective in *ompR* and *bolA* were refractile to the addition of exogenous cAMP with regard to cell width (Figs 6E and 6F).

### Defects in acetate metabolism support links between fatty acid synthesis and assembly of the cell division machinery

Cell cycle progression is tightly correlated with growth rate in wild-type cells. Fast growing cells are not only larger but contain a higher frequency of FtsZ rings (Z-rings) with approximately 90% of *E. coli* cells containing visible Z-rings in LB-glucose. Meanwhile, slow growing cells contain fewer Z-rings (∼50%). Z-ring frequency shows a roughly linear relationship to growth rate when grown in different media (Fig 4C dotted line) [12,39]. Taking advantage of a green fluorescent protein (GFP)-FtsZ fusion that allows us to label the FtsZ ring without detectably altering size or growth [6], we assessed the frequency of FtsZ ring formation in the 17 mutants in Classes I through V.

Although the Z-ring frequency and growth rate were correlated in the majority of CCM mutants, several exhibited a disconnect between these two parameters, specifically those associated with acetate metabolism (black squares in Fig 4C). Cells defective in *aceE,* the E1 protein of pyruvate dehydrogenase that converts pyruvate to acetyl-CoA, exhibited an extremely high (86%) Z-ring frequency relative to wild type cells cultured in medium of the same growth rate (∼75%) [12], while both *ackA* and *pta,* involved in converting acetyl-CoA to acetate, exhibited lower Z-ring frequencies relative to growth rate (69% and 67% respectively). Assembly of the cell division machinery is a multistep process involving first, recruitment of FtsZ and other so-called early proteins to the nascent division site followed by a brief delay and assembly of the late proteins, which include enzymes required for synthesis of the septal cell wall (reviewed in [40]). In *E. coli*, division is coincident with recruitment of FtsN, the last protein to be incorporated into the division machinery and the putative “trigger” for cross wall synthesis [41,42]. The two-step nature of division machinery assembly suggests two potential explanations for the elevated Z-ring frequency in *aceE::kan* mutants: 1) premature recruitment of FtsZ to the septum or 2) delays in late protein recruitment and activation of cytokinesis. Conversely, the lower frequency of FtsZ rings in *ackA::kan* and *pta::kan* mutants could be a consequence of either 1) delays in FtsZ recruitment or 2) premature late protein recruitment and/or activation of division mediated cell wall synthesis.

To clarify the underlying cause of the elevated Z-ring frequency in *aceE::kan* cells and depressed Z ring frequency in the *ackA::kan* and *pta::kan* backgrounds we took advantage of a heterologous, IPTG-inducible FtsN-GFP [43] to assess the efficiency of FtsN recruitment in all three mutants. Growth in LB-glu showed 32% of wild-type cells with FtsN-GFP localization at mid-cell (Fig 7A). In contrast, despite varying Z-ring frequencies, all three mutants exhibit more or less the same frequency of FtsN ring formation (24%, 23%, and 26% for *aceE::kan,ackA::kan*, and *pta::kan* respectively) with all three being significantly less than wild type. The consistency of FtsN ring frequency, despite differences in Z-ring frequency between the three CCM mutants, suggests that it is delayed recruitment of the late division proteins and/or activation of cytokinesis that is responsible for the high Z-ring frequency in *aceE::kan* cells while the lower Z-ring frequeny in *ackA::kan* and *pta::kan* mutants would appear to be a consequence of a delay in FtsZ assembly.

Acetyl-CoA metabolism plays a critical role in the ability of cells to synthesize the wide variety of fatty acids and lipids that constitute the cell envelope [44]. *aceE*, in particular, encodes a subunit of the pyruvate dehydrogenase complex responsible for aerobic synthesis of the majority of acetyl-CoA, a necessary precursor in the first step of FAS [45]. Defects in aceE are thus expected to reduce flux through FAS, and based on previous work, shift the balance in lipid composition [46]. Although not directly involved in FAS, defects in pta and *ackA* may lead to a build-up of acetyl-CoA and potentially also shift flux through FAS.

Based on previous studies linking lipid composition to cell morphology [47], we speculated defects in acetate metabolism might result in changes in plasma membrane composition that interfere with assembly and activity of the cell division machinery. To test this idea, we measured the Z-ring frequency of wild type and mutant cells cultured in the presence of sub-inhibitory concentrations of the fatty acid synthesis inhibitor triclosan (50 ng/mL). This concentration is too low to impact growth rate but has a modest positive impact on cell size [6]. Triclosan inhibits FabI, the enzyme that catalyzes the reduction of enoyl-ACP during the elongation cycle of fatty acid synthesis [48]. Although not a perfect recapitulation of a block upstream of acetyl-CoA synthesis, as it blocks a downstream step, both triclosan treatment and the aceE mutations should similarly curtail flux through FAS.

**Figure 7:**
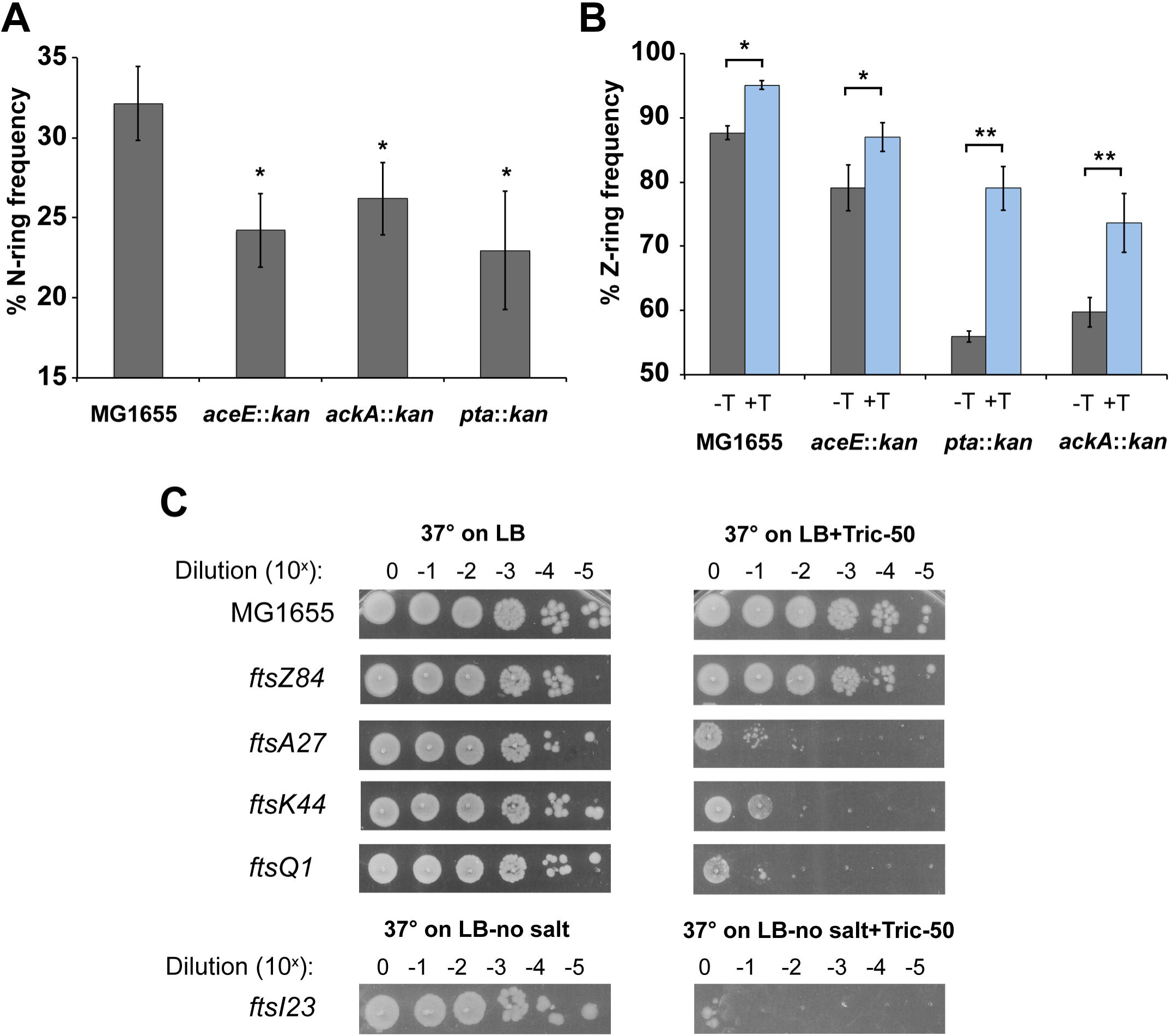
Partial inhibition of fatty acid synthesis negatively impacts assembly and activity of the late division proteins. (A). FtsN septal localization was measured in MG1655, *aceE::kan, ackA::kan*, and *pta::kan* expressing an *ftsN-GFP* from an IPTG inducible promoter (B.) FtsZ-ring frequency was measured MG1655, aceE::kan, ackA::kan, and pta::kan cells expressing ftsZ-GFP from an IPTG-inducible promoter. Cells were cultured with (gray) or without (black) 50 ng/mL triclosan. (C) Representative plating efficiencies of wild-type cells and the heat-sensitive cell division mutants *ftsZ84, ftsA27, ftsK44,* and *ftsQ1* cultured on LB at 37°C with or without triclosan. *ftsI23* was plated on LB-no salt at 37°C with or without triclosan.

In support of a model in which modest reductions in flux through FAS negatively impact assembly of the cell division machinery, Z-ring frequency was significantly increased in wild type cells as well as in *aceE::kan, pta*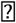*kan*, and *ackA*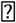*kan* cells during growth in sub-MIC (minimum inhibitory concentration), 50 ng/mL triclosan. (Z-ring frequency: wild type 88% +T/95%-T; pta::kan 56%-T/ 79% + T; *ackA::kan* 60%-T/74% +T; *aceE*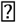*kan* 79%-T to 87% +T) (Fig 7B). *pta* or *ackA* deletion mutants were slightly more sensitive to triclosan with regard to Z-ring frequency than either *aceE* or wild type cells, likely a reflection of the already high frequency of FtsZ rings in these latter strains.

Further supporting a positive connection between FAS and assembly of the cell division machinery, 50 ng/mL triclosan enhanced the heat sensitive phenotype of conditional mutations in several cell division genes including *ftsA (ftsA27*-an early, actin-like protein), *ftsK (ftsK44*—a late division protein involved in chromosome partitioning), *ftsQ (ftsQ1*—a late division protein believed to serve primarily as a scaffold), and *ftsI (ftsI23*—a late division protein with a trans-peptidase domain) [49–52]. Somewhat surprisingly given its impact on FtsZ ring frequency (Fig 7B), triclosan had no impact on the heat-sensitivity of *ftsZ84* (Fig 7C). We speculate that the differential impact of triclosan on ftsZ84 relative to the other conditional mutants is a consequence of their subcellular location (cytoplasmic – FtsZ) versus membrane associated (FtsA) and integrated into the membrane (FtsQ, FtsK, and FtsI) [40].

## Discussion

Our systematic analysis of the relationship between central carbon metabolism, cell growth, and cell size both reinforces the Growth Law as a general principal (i.e. growth rate and size are positively correlated) but also calls into question many of the fundamental assumptions that underlie it. While the largest class of mutants with observable phenotypes—Class II: small with reduced growth rates, essentially “obeyed” the law so to speak, growth rate and size varied independently in the other classes of CCM mutants (Fig 1B). Class II mutants closely resemble cells cultured in nutrient-poor conditions with regard to growth rate and size, suggesting these cells are experiencing a near global reduction in biosynthetic capacity. In contrast, growth rate and morphology are independent of one another in the other nine CCM mutants (Table 1), consistent with defects in specific regulatory or structural functions that directly impact cell length and/or width.

These results are in line with accumulating work indicating that growth rate and cell size are independent phenomena. Modest defects in cell division are well known to increase bacterial cell size—particularly length—without impacting growth rate [4,7,50,51,53], while defects in transcription severely curtail growth but have little detectable impact on morphology [6]. Most importantly, an exhaustive phenotypic analysis of the entirety of non-essential genes in *E. coli* by the Jacobs-Wagner group, failed to identify any evidence supporting a direct connection between growth rate and size [54].

### Overflow metabolism and cell size

In a common, yet paradoxical, fermentative mechanism known as “overflow metabolism”, *E. coli* convert up to 50% of available glucose into acetate during aerobic growth in nutrient rich conditions [55]. Acetate fermentation is inefficient, resulting in ∼14 fewer ATP per glucose than oxidative phosphorylation and, unlike other fermentative pathways, does not regenerate NAD^+^ from NADH [56]. While it is unclear why bacteria perform overflow metabolism, it is potentially a more proteome-efficient means of moving carbon through CCM, ensuring that glycolysis proceeds at a maximum pace when nutrients are plentiful.

Supporting the idea that overflow metabolism is critical for maximizing glycolytic flux, defects in *pta, ackA*, and *aceE*, encoding the first two enzymes in acetate fermentation and a subunit of pyruvate dehydrogenase respectively, are dramatically impaired in both growth rate and cell size. This is consistent with a global reduction in biosynthetic capacity. Whether this reduction is the result of lower ATP production or the consequence of a bottleneck in flux through critical biosynthetic pathways is unclear. Regardless, we suspect that reductions in ATP levels and/or biosynthesis indirectly result in accumulation of the alarmone ppGpp [57], providing an explanation for the observed suppression of *ftsZ84* heat-sensitivity in *pta, ackA*, and *aceE* deletion strains. Increases in ppGpp are well known to suppress the conditional nature of *ftsZ84*, although the precise mechanism has yet to be determined [17,58].

### cAMP is a key player in maintaining the correct cellular aspect ratio

A somewhat surprising finding is the identification of cAMP as a contributor to nutrient dependent reduction in both cell length and width. Accumulating under gluconeogenic conditions, cAMP binds to the transcriptional activator Crp and regulates expression of more the than 280 genes [59]. Although the precise member(s) of the cAMP-Crp regulon responsible remains elusive, our data indicate that cAMP-Crp function together to reduce width under nutrient poor conditions via their impact on the morphogenic factor BolA and cell length via its impact on the assembly of FtsZ (Figs 5 and 6). In this regard, cAMP differs from metabolic signaling molecule, ppGpp, which genetics indicate is a positive regulator of division (Fig 8).

**Figure 8:**
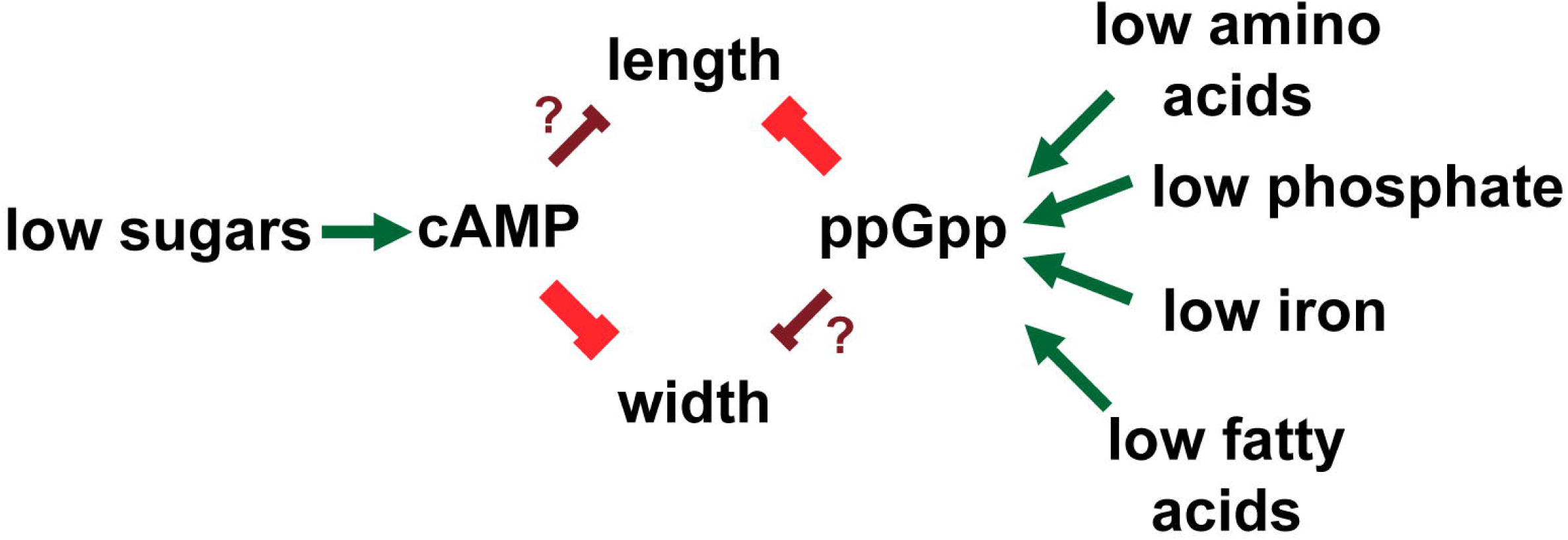
Proposed pathway linking nutrient levels to cell length and width through the signaling nucleotides cAMP and ppGpp.

### CCM is robust but not necessarily efficient

Perhaps the biggest surprise of this study is our finding that the defects in the majority of non-essential CCM genes had little to no detectable impact on either cell size or growth rate. While the lack of phenotype in many cases is likely a consequence of our screening conditions, nutrient-rich, complex medium supplemented with glucose, in certain cases the robust growth we observed is difficult to explain. For example, we anticipated that defects in phosphoglucose isomerase (encoded by *pgi)*, the first dedicated step in glycolysis would dramatically impact growth rate and size by virtue of its impact on not only glycolysis, but also other associated pathways leading to reductions in ATP synthesis and overall biosynthetic capacity [60]. Instead, we were unable to distinguish *pgi* mutants from wild type cells during growth LB-glucose (Table S2). Based on our identification of seven mutants in the pentose-phosphate pathway, we hypothesize that despite the ubiquitous nature of glycolysis in nutrient- rich medium, glucose is predominantly driven through the pentose-phosphate pathway to generate essential building blocks.

Similarly, we observed little in the way of a detectable phenotype associated with defects in the gene encoding citrate synthase *gltA.* Essential for the complete oxidation of carbon, GltA synthesizes citrate from oxaloacetate and acetyl-CoA at the entry point into the TCA cycle [61]. Despite its seeming importance, previous work indicates that *E. coli* represses *gltA* transcription in the presence of glucose, suggesting that it can forgo complete carbon oxidation under nutrient rich conditions [59,62]. It may be that under nutrient rich condition, *E. coli* maximizes rapid growth over the efficient use of available carbon, a proposition that is in line with our data indicating that defects in acetate metabolism significantly curtail growth in nutrient rich medium (Table 1).

### Concluding thoughts

While this screen demonstrates the ability of systematic phenotypic analysis to identify new connections between CCM, cell growth and cell size, many questions remain unanswered. In addition to the large number of insertion mutants with no observable phenotype, defects in 17 genes resulted in dramatic changes in growth rate and cell morphology. Of these, we were able to identify the proximal cause in 24% (*crr, aceE, pta, ackA*; Figs 5–7). A better understanding of the remaining 13 mutations will likely require a combination of genetics and metabolomics to tease apart the root cause of their morphological and growth related defects.

Additionally, although this effort focused specifically on carbon metabolism, the availability of other nutrients—particularly nitrogen, phosphorous, sulfur and iron— have also been implicated in growth and morphology. Assessing the phenotype of the CCM insertion collection under conditions in which these factors are limiting is thus also worthwhile. Finally, *E. coli* is believed to spend much of its time under microaerobic or anaerobic conditions. Understanding how oxygen levels impact growth, cell cycle progression, and cell morphology thus remains a tantalizingly open question.

## Materials and Methods

### Strains and media

All chemicals, media components, and antibiotics were purchased from Sigma-Aldrich (St. Louis, MO). *E. coli* strains were constructed through standard P1 vir transductions [63] moving kanamycin marked insertion deletions from Keio collection strains [18] purchased from the Coli Genomic Stock Center into MG1655 (*F*^-^ *λ*^-^ *ilvGrfb*-50 *rph*-1), our “wild type” parental strain. Cell were cultured in Luria-Bertani (LB) broth supplemented with 0.2% glucose or AB defined media [64] supplemented with either 0.2% glucose or 0.2% succinate as the sole carbon source. Unless otherwise noted (e.g. for heat sensitive conditional alleles of cell division genes including *ftsZ84, ftsA27, ftsK44, ftsQ1*, and *ftsI23*), cells were cultured at 37° C and sampled for analysis at OD_600_ between 0.1 and 0.2. For the primary analysis cells were cultured in 5 mL LB + 0.2% glucose and aerated at 200 rpm. For measuring the effect of different media, cells were cultured in 200 μL of specified media in a 96-well plate and aerated at 567 rpm in a BioTek^©^Eon^©^ plate reader (Fig 3).

### Microscopy

Cells were sampled directly from cultures and fixed in 16% paraformaldehyde (Electron Microscopy Sciences) as described [6]. Fixed cells were mounted on agarose pads and visualized using a 100x phase-fluor objective on a Nikon^©^ Ti-E inverted microscope. To facilitate visualization of FtsZ rings, 1 mM IPTG was added to cultures of cells encoding a chromosomal P_IPTG_-ftsZ-GFP [65]. Expression of a chromosomal P_IPTG_-ftsN-GFP was induced with 50 µM IPTG [43]. GFP fluorescence was detected with a C-FL GFP filter cube with 470/40nm excitation and 525/50nm emission (Chroma Technology Corporation) Nikon Elements software (Nikon Instruments, Inc.) was used for image capture and analysis. >200 cells were counted per experimental replicate and at least three independent replicates were used for each data point.

### Cell Size Measurements

Cell size from phase-contrast images was determined with one of two freely available segmentation software packages: the ImageJ^©^ plug-in Coli Inspector [19] (Fig 1,Tables 1 and S2) or SuperSegger, a MatLab^©^ based application [20] (Figs 3–7, Table S3). As Coli Inspector does not determine area directly, we estimated it as length x width. SuperSegger directly determines the area of individual cells. We determined the area of ≥100 cells for each experiment and/or condition. Unless otherwise noted, average size was determined from 3 independent experiments.

### Dot plating efficiency assays

Heat-sensitive cell division alleles, *ftsZ84, ftsA27, ftsK44, ftsQ1*, and *ftsI23* were cultured in LB at 30°C degrees to an OD_600_ of ∼0.5. 1 mL of cells were then pelleted, washed once with LB-no salt, and re-suspended to an OD_600_ of 0.1 in LB-no salt. 10 μL of serial dilutions between 10^-1^ to 10^-6^ were plated under permissive conditions (LB at 30°C) or non-permissive conditions (LB-no salt at 37°C). Due to the strains extreme sensitivity, LB-no salt at 30°C was non-permissive for *zwf::kan ftsZ84* double mutants. For cAMP or triclosan addition experiments, 5 mM cAMP or 50 ng/mL triclosan was added directly to LB agar prior to pouring the plates. Each plating efficiency experiment was repeated three times with representative images shown.

## Acknowledgements

We would like to thank the Coli Genetic Stock Center (CGSC) for providing numerous strains quickly. We’d like to thank Norbert Vischer for aid in using Coli Inspector and Stella Stylianidou and Paul Wiggins for help with SuperSegger. We would like to thank Elizabeth Mueller for work on the *pgm ftsZ84* double mutant. We would also like to thank Elizabeth Mueller, Dr. Ashley Sherp, and the Hani Zaher laboratory for critical feedback on this work.

## Supporting Information

**Table S1. List of bacterial strains used in this study**

**Table S2. Raw size and growth rate data for all central carbon metabolic gene deletions.**

**Table S3. Raw data for figure 3 (CCM mutant growth in different media).**

**Table S4. Raw data for figure 4C (Z-ring frequency of CCM mutants).**

